# Comprehensive long-read transcriptome analysis uncovers alternative RNA processing feature and isoform diversity in ovarian cancer progression

**DOI:** 10.64898/2026.03.11.710989

**Authors:** Tianxiang Liu, Jianying Lv, Shuo Wang, Yanhua Liu, Yanan Chen, Jia Li, Longlong Wang, Yi Shi, Wei Ding, Yongjun Piao

**Author notes:** Corresponding author. (Y.P.); (W.D.). These authors contributed equally to this work (Tianxiang Liu, Jianying Lv, Shuo Wang).

## Abstract

Post-transcriptional processing has a crucial yet largely unresolved dynamic change and role during the malignant progression of ovarian cancer, especially due to the limited read length of short-read RNA sequencing being insufficient to capture transcript diversity. Here, we performed Iso and RNA sequencing on paired normal, primary tumor, and metastatic samples, generating a comprehensive isoform atlas of over 41,000 full-length transcripts including many unannotated isoforms. Integrative analyses revealed extensive isoform-level remodeling across disease states that often occurred without concordant alterations at the gene level, emphasizing the importance of qualitative transcript regulation. Notably, we identified isoform-level alterations with distinct biological and clinical relevance, including differential expression of the short KRAS isoform, a tumor-specific isoform switch of TMEM201, and an alternative first-exon event in FNDC3B associated with poor survival. Together, these findings provide a high-resolution map of the ovarian cancer transcriptome and illustrate how long-read sequencing exposes multiple layers of post-transcriptional and clinical insight that remain hidden in conventional expression profiling.

## Introduction

Ovarian cancer is one of the most common gynecologic malignancies[1]. Recent global cancer statistics indicated that an estimated 324,398 women were newly diagnosed and 206,839 deaths occurred in 2022[2]. Despite advances in screening methods and therapeutic strategies, the prognosis of ovarian cancer remains poor, largely because most cases are detected at advanced stages and are prone to recurrence and metastasis[3]. Therefore, understanding the molecular events that drive tumorigenesis and dissemination is crucial for improving clinical outcomes.

Transcriptome-level alterations, such as dysregulated gene expression[4], aberrant alternative splicing[5, 6], and changes in polyadenylation[7], are key contributors to cancer initiation and progression. Conventional RNA sequencing (RNA-seq) has provided extensive information on transcriptional regulation across cancers. This technology delivers accurate base calling (>99.9%), supports parallel analysis of multiple samples, and has been widely adopted for transcriptomic profiling. Nevertheless, the limited read length constrains its ability to reconstruct complete isoforms and resolve complex splicing patterns[8]. As a result, our understanding of transcriptomic heterogeneity in ovarian cancer remains incomplete.

More recently, long-read sequencing (LR-seq) technologies have provided a promising solution by generating transcript-length reads, typically ranging from 1 to 100 kb[9]. It allows direct identification of full-length isoforms, discovery of previously unannotated transcripts[10], and precise analysis of alternative splicing[11, 12] and polyadenylation[13]. Long-read platforms have already been applied in diverse contexts, including the study of promoter usage in gastric cancer subtypes[14], discovery of isoform biomarkers in primary and metastatic liver cancer[15], and exploration of region-specific transcriptional patterns in the human brain samples[16]. Here, we applied LR-seq to nine tissue samples obtained from three ovarian cancer patients, including matched normal ovarian tissues, primary tumors, and metastatic lesions. By comparing these three disease states within the same individuals, we aimed to uncover transcriptomic alterations accompanying tumor initiation and metastasis. Our analyses revealed isoform-level changes involving splicing and polyadenylation that may contribute to metastatic potential. These findings shed light on molecular features of ovarian cancer progression and underscore the value of long-read transcriptomics in cancer research.

## Results

### Isoform landscape in normal, primary, and metastatic ovarian cancer tissues

We collected nine tissues from three patients with high-grade serous ovarian cancer (HGSOC), including normal tissues, primary ovarian tumors, and omental metastases. These samples were sequenced using Iso-Seq long-read and DNBSEQ-T7 RNA-seq platform to comprehensively investigate the transcriptome of HGSOC (Fig. 1A). LR-seq yielded an average of 6 million non-chimeric reads per sample, defined as sequences spanning from the 5’ to 3’ ends and including the poly(A) tail (Table S1). Following clustering and polishing, each sample produced approximately 430,000 circular consensus sequencing (CCS) reads on average, resulting in a total of 4,000,000 reads for downstream analysis. These CCS reads had an average length of 1.6 kbp, with over 99% successfully mapped to the reference genome[17]. Isoforms were detected and quantified using IsoQuant[18], a tool recommended by the LR-seq Genome Annotation Assessment Project (LRGASP) for transcriptome analysis of well-annotated organisms[19]. We then categorized the detected isoforms into five distinct groups in each sample: Full Splice Match (FSM), in which splice junctions exactly matched the reference; Incomplete Splice Match (ISM), in which only a subset of the consecutive splice junctions match the reference; Novel Not in Catalog (NNIC), which contains novel splice junctions; Novel in Catalog (NIC), where the splice junctions are known but the isoform is novel, and Others, for transcripts that do not fit into any of these groups (Fig. 1B). The majority of detected isoforms were FSM, followed by ISM and NIC, with NNIC isoforms representing a smaller proportion, and this distribution was consistent across normal tissues, primary tumors, and metastases.

**Fig. 1.**
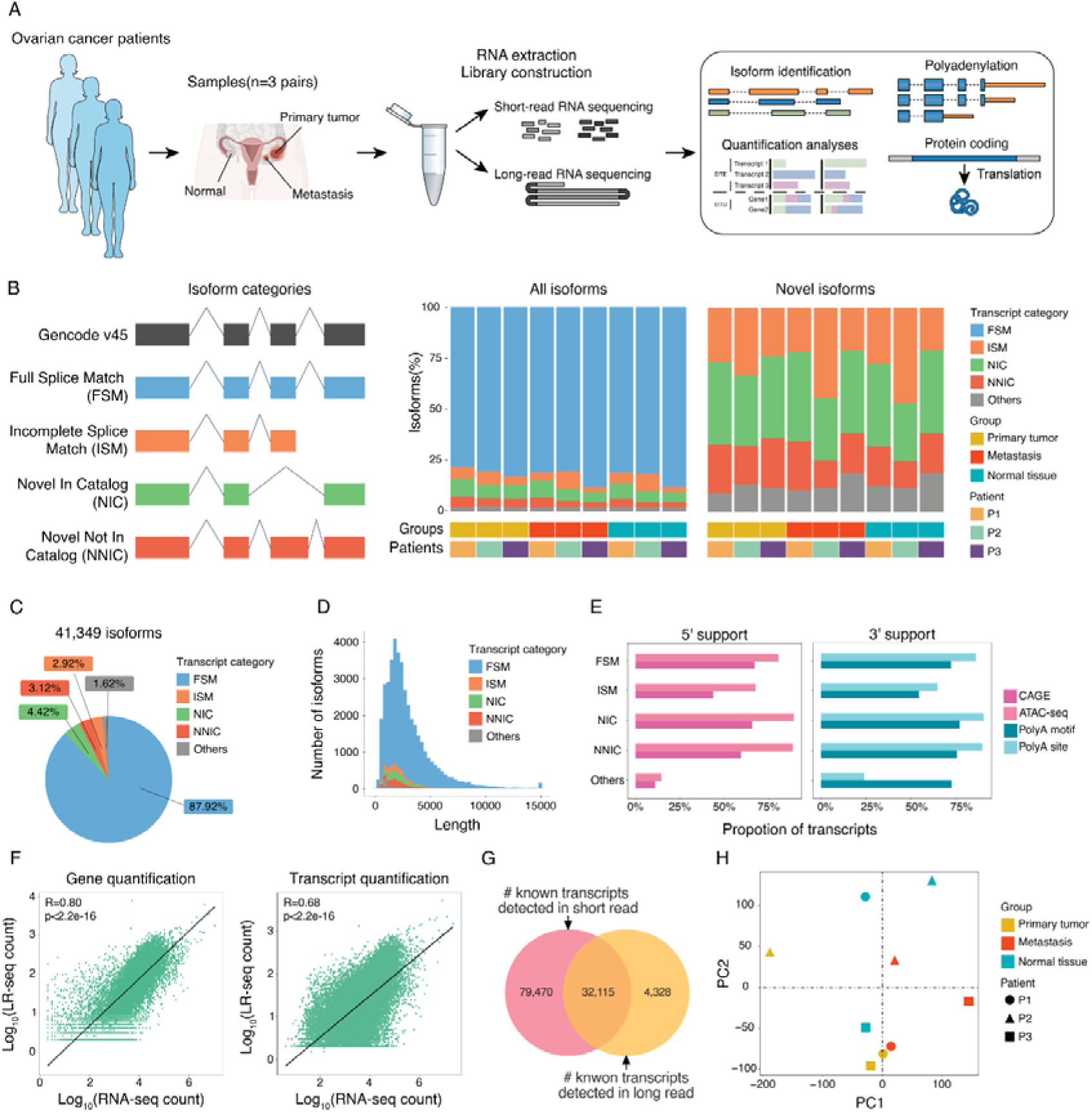
Full-length transcriptome profiling of high-grade serous ovarian cancer (HGSOC) patients using Iso-seq. (A) Schematic overview of the study design. Matched normal, primary, and metastatic samples from three HGSOC patients were subjected to both long-read (Iso-Seq) and short-read (DNBSEQ-T7) RNA sequencing to determine transcript dynamics. (B) Classification of isoforms into five transcript categories, including Full Splice Match (FSM), Incomplete Splice Match (ISM), Novel in Catalog (NIC), Novel Not in Catalog (NNIC), and others, based on splice-junction concordance with GENCODE v45. Bar plots show the relative proportions of each category across patients and tissue types. (C) Distribution of the total isoforms identified in the HGSOC transcriptome. (D) Isoform length distribution for each transcript category. (E) Proportion of transcripts supported by multi-omic datasets. (F) Correlation between long-read and short-read quantification at the gene and transcript levels. (G) Venn diagram showing the overlap of isoforms detected by short-read and long-read sequencing. (H) Principal component analysis (PCA) of isoform expression profiles.

In total, 41,349 distinct isoforms were identified in our HGSOC transcriptome, corresponding to approximately 20,000 genes. Our analysis revealed that approximately half of the genes were consistently detected in all samples, and at least two-thirds were shared among the three groups (Fig. S1A). Of all genes annotated in GENGODE v45, ∼75% of protein-coding genes were identified, most of which displayed high expression levels. In contrast, only a limited number of lowly expressed lncRNA genes were detected, while pseudogenes were rarely observed (Fig. S1B). The vast majority of these isoforms (87.92%) were FSM, with smaller proportions of ISM (3.12%), NIC (4.42%), and NNIC (2.92%) isoforms (Fig. 1C).

Isoform-level analysis showed a higher degree of sample specificity, with ∼17% of isoforms detected exclusively in a single sample (Fig. S1C). FSM isoforms accounted for the largest proportion of sample-specific transcripts, whereas novel NIC and NNIC isoforms were detected across most samples (Fig. S1D), Most of the novel isoforms were distributed across all three groups (Fig. S1E), suggesting that IsoQuant employs a strict filtering strategy for novel isoform identification. The length distribution of the isoforms revealed that the majority ranged from 1 to 2 kb, with a smaller population of longer isoforms extending up to approximately 15 kb (Fig. 1D), Meanwhile, most isoforms contained fewer than 20 exons (Fig. S1F). No significant differences in isoform length were observed across different transcript categories. To assess the reliability of the detected isoforms, we analyzed the support for 5’ and 3’ ends of the transcripts using multi-omics datasets. More than 75% of the 5’ ends of FSM, NIC, and NNIC transcripts overlapped with transcription start sites (TSSs) identified by CAGE or open chromatin regions from the TCGA pan-cancer cohort. Similarly, the 3’ ends of FSM, NIC, and NNIC transcripts were supported by poly(A) sites validated by 3’-seq or the canonical poly(A) motif (Fig. 1E). In contrast, ISM isoforms had less end support, likely due to absence of first or last exons, which may limit the detection of their 5’ and 3’ ends. Overall, these results highlighted that both known and novel isoforms detected in our study have broad support from different experimental techniques.

Comparative analysis indicated that expression quantification from LR-seq exhibited a more concentrated distribution at both the gene and isoform levels, while showing lower overall expression values compared with short-read sequencing (Fig. S1G). We examined the correlation of gene and isoform expression levels from LR-seq and RNA-seq, respectively. Strong correlations were observed (Fig. 1F), with gene-level quantification showing a correlation of 0.80 (*p*-value < 2.2e-16) and transcript-level quantification showing a correlation of 0.68 (*p*-value < 2.2e-16). However, the number of isoforms detected by the two platforms differed dramatically, with 111,585 isoforms detected by RNA-seq and 36,443 detected by Iso-seq (Fig. 1G). Of these, 32,115 isoforms overlapped between the two platforms, a relatively small number for short-read sequencing but a significant proportion for LR-seq. This discrepancy suggests that short-read sequencing may result in a large number of false positives. This is likely due to the limited read length of RNA-seq, which requires the assembly of short reads using computational techniques, potentially leading to the generation of incorrect isoforms. Additionally, principal component analysis was performed to assess the ability of transcripts to distinguish normal tissues, primary tumors, and metastases. The results revealed that primary tumor and metastasis samples clustered separately, with each patient’s samples exhibiting distinct patterns. Notably, normal tissue samples formed a separate cluster from both primary tumors and metastases, indicating significant transcriptomic differences between these tissue types (Fig. 1H). Taken together, these findings provide an in-depth view of the full-length transcriptome landscape in HGSOC and highlights the value of LR-seq for uncovering novel isoforms with clinical significance.

### Functional and proteomic characterization of isoforms detected by LR-seq

We examined the coding potential of the isoforms detected through LR-seq by analyzing the open reading frames (ORFs) and their alignment to Uniprot databases[20]. To examine how isoform diversity translated into protein-level variation, we extracted the best ORF for each isoform. Overall, most isoforms from all categories were predicted to have coding potential based on the protein prediction (Fig. S2A). Approximately 90% of FSM, NIC, and NNIC isoforms were predicted with ORFs, while around 85% of others like ISM and intergenic isoforms had coding regions. However, we observed that 12% of NIC and 17% of NNIC transcripts contain premature termination codons (PTCs) within their ORF regions, making them more likely to undergo mRNA degradation through the nonsense-mediated decay (NMD) pathway during translation, while only 6% of FSM ORFs contain PTCs (Fig. S2B).

To distinguish novel ORFs from known ORFs, the predicted ORFs from LR-seq were aligned to the most similar sequences in the UniProt database (Fig. 2A). Nearly 70% of ORFs from FSM shared over 99% identity with the matched UniProt sequences. Conversely, less than half of ISM (25%), NIC (37%), NNIC (37%) LR-seq isoform encode ORFs were matched in UniProt. Therefore, alterations in isoform structure give rise to novel proteomic diversity in ovarian cancer. We further investigated the NMD mechanism for novel ORFs and found that the proportion of PTC-containing novel ORFs in NIC and NNIC isoforms was comparable to that observed across all ORFs. By contrast, FSM isoforms exhibited a higher probability of mRNA degradation through NMD (Fig. S2B). This suggests that alterations in ORF sequences may introduce PTCs, leading to an increased occurrence of nonsense mutations.

**Fig. 2.**
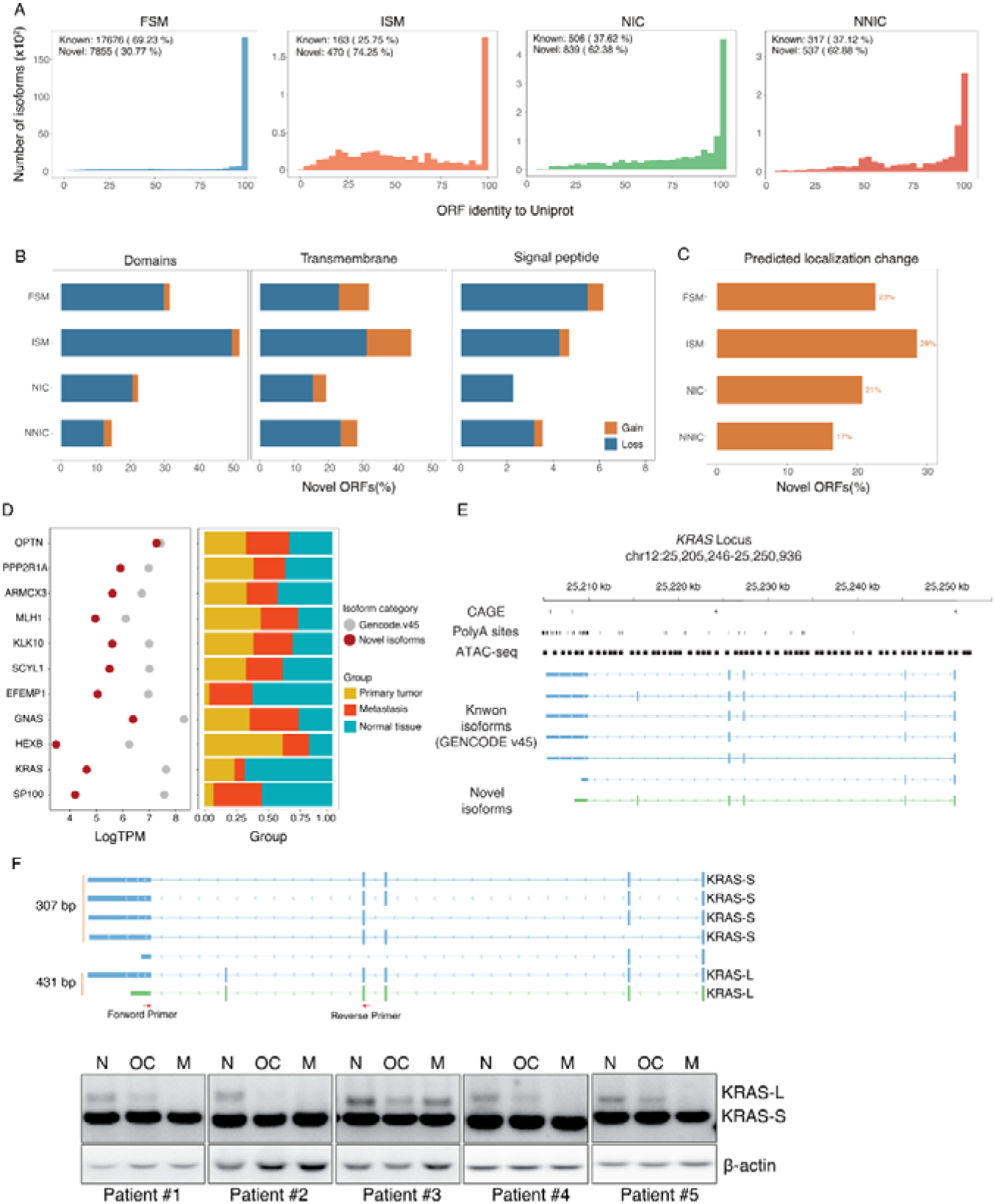
Characterization of potential functional consequences of HGSOC transcriptome at protein level. (A) Distribution of open reading frame (ORF) identity between long-read–derived isoforms and their best matches in the UniProt database across transcript categories. (B Functional features gained or lost in novel ORFs, including protein domains, transmembrane regions, and signal peptides. (C) Proportion of subcellular localization changes of novel ORFs compared with reference proteins. (D) Representative genes showing expression of novel isoforms in normal, primary, and metastatic ovarian cancer tissues. (E) Example of the KRAS locus illustrating integration of CAGE, ATAC-seq, and poly(A)-site data. (F) Experimental validation of novel KRAS isoforms in five patient pairs.

Then we explored the potential impact of post-transcriptional processing on the functional changes of novel ORFs. We first predicted transmembrane regions of the ORF sequences and searched for protein domains in the Pfam[21] database (Fig. 2B). Approximately 20% of novel ORFs exhibited loss of transmembrane regions or domains, while a small fraction showed gains of either feature, suggesting potential modifications in protein structure and function. Prediction of signal peptide regions revealed that most (90%) ORFs lacked signal peptide sequences. Notably, ∼5% of novel ORFs displayed loss of signal peptides, accounting for nearly half of those that originally contained such sequences (Fig. 2B). This loss is likely to affect protein function. In addition to influencing protein function, modifications in protein domains can alter subcellular localization, which can be predicted using DeepLoc[22] (Fig. 2C). Approximately 20% of novel ORFs exhibited changes in subcellular localization, with the most frequent alterations occurring between the cytoplasm and nucleus, followed by changes between the cytoplasm and mitochondria (Fig. S2C). Proteomic validation of novel ORFs identified by LR-seq was performed using CPTAC datasets[23], including five cohorts with 1,344 samples from 440 patients. The reliability of these ORFs was evaluated by scoring tandem mass spectrometry (MS/MS) spectra against theoretical peptides generated from sequencing data. Notably, 80% of novel ORFs were matched in the proteomic datasets, providing clinical support for their existence, although they remain unannotated in UniProt (Fig. S2D).

Interestingly, although KRAS is well known as an oncogene, we observed that its overall expression level was elevated in normal ovarian tissues (Fig. 2D). To explore this unexpected observation, we searched public transcriptomic databases and identified multiple KRAS transcript isoforms (Fig. 2E). Based on LR-seq data, we confirmed the presence of two major isoforms, termed KRAS-L and KRAS-S. Isoform-specific primers were designed to validate their expression patterns in normal and tumor ovarian tissues. Consistent with our sequencing results, KRAS-L was predominantly expressed in normal ovarian tissues (Fig. 2F). This finding is also in agreement with previous reports describing isoform-specific expression of KRAS in different tissue contexts[24].

### Isoform switching during ovarian cancer progression

We next conducted comparative analyses at multiple transcriptional levels, including gene expression, transcript abundance, and isoform usage, in normal, primary, and metastatic tissues. As illustrated in Fig. 3A, differential gene expression (DGE) and differential transcript expression (DTE) analysis determined significantly expressed genes and transcripts between conditions while differential transcript usage (DTU) examines the expression changes of multiple transcripts belonging to same gene between conditions (Table S2). In total, we identified 1,134 genes that showed both significant DGE and DTE when comparing cancer (including primary and metastatic tumors) to normal tissues, and 40 genes demonstrated simultaneous DTU (Fig. 3B, left panel). Only a subset of genes with changes in overall expression overlapped with those exhibiting transcript-level or isoform usage alterations, highlighting the complexity of post-transcriptional process in HGSOC. Comparisons between metastatic and primary tumor samples indicated fewer significant changes, with 222 genes exhibiting both DGE and DTE, and 130 genes demonstrating DTU only (Fig. 3B, right panel). These results indicate extensive isoform-level transcriptional reprogramming during ovarian tumorigenesis, which appears to relatively less vary in metastatic progression. Functional analysis of differential expression patterns between cancer and normal tissues revealed that DGE- and DTE-associated genes were enriched in cell cycle and DNA repair pathways, including G2/M checkpoint, E2F targets, and UV response signatures (Fig. 3C, left panel). In metastatic tumors, functional enrichment shifted toward metabolic adaptation, immune modulation, and post-transcriptional regulation such as nonsense-mediated decay (Fig. 3C, right panel).

**Fig. 3.**
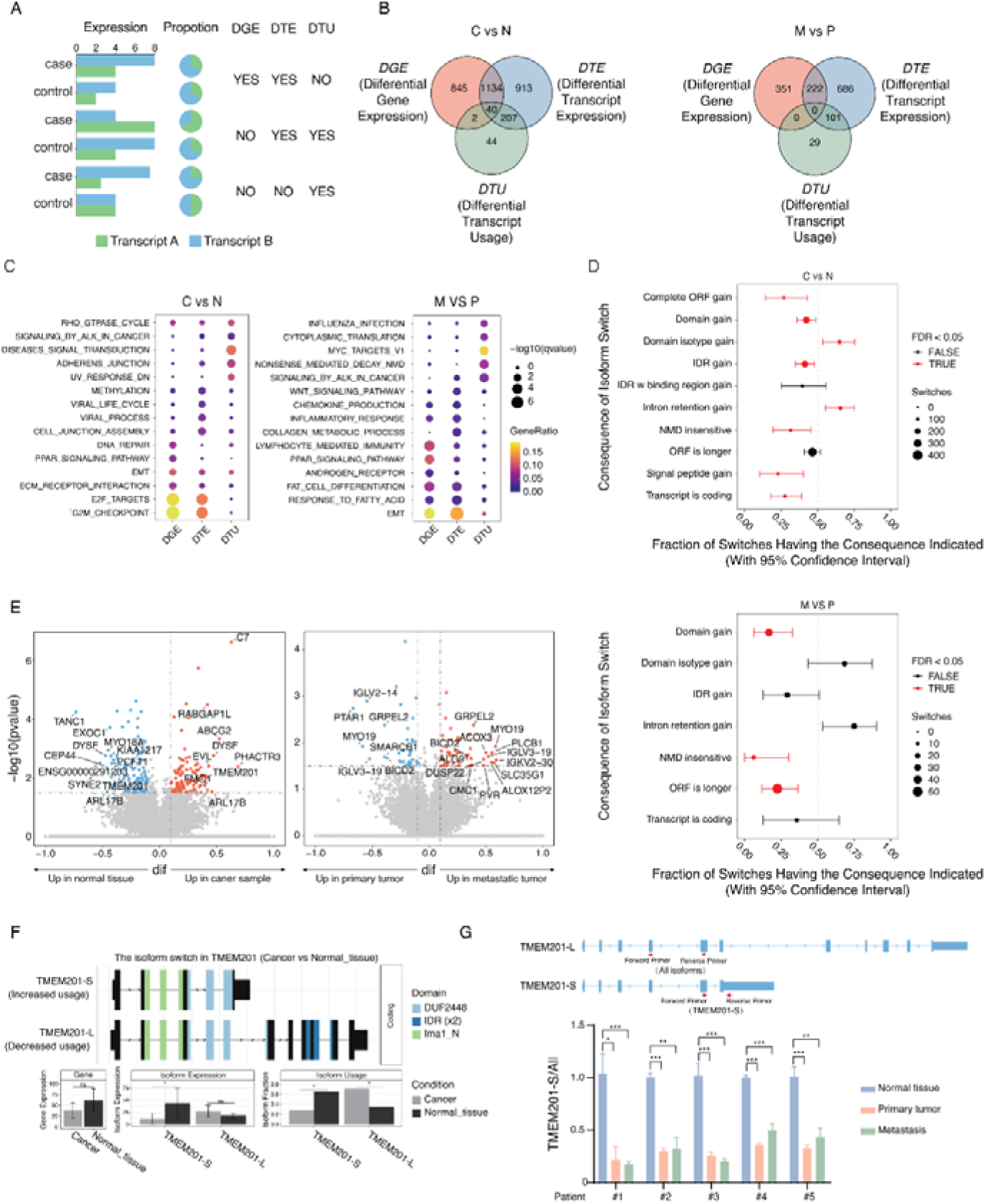
Determination of isoform switches in HGSOC. (A) Illustration of differential gene expression (DGE), differential transcript expression (DTE), and differential transcript usage analysis (DTU). (B) Venn diagrams showing overlap among DGE-, DTE-, and DTU-identified genes. (C) Enrichment analysis of genes with significant DGE, DTE, or DTU. (D) Predicted functional consequences of isoform switches between groups. (E) Volcano plots of genes with significant DTU between cancer and normal samples (left) as well as between metastatic and primary tumors (right). (F) Illustration of structure and usage of TMEM201 isoforms. (G) RT-PCR validation of TMEM201 isoform switching across five patient pairs.

To understand the biological relevance of these isoform-level changes, we predicted the functional consequences of isoform switches between conditions (Fig. 3D). The isoform switches identified between normal and cancer tissues frequently involved gains of complete ORF and intrinsically disordered regions (IDRs). In addition, comparisons between metastatic and primary tumors showed fewer switching events with significant consequences. Nevertheless, notable isoform-level alterations were observed, including gains of novel domains, introduction of insensitive NMD or longer ORF, highlighting ongoing transcriptomic remodeling in metastatic progression. Several candidate genes displayed distinct patterns of isoform usage, suggesting their potential as isoform-level biomarkers for distinguishing normal, primary, and metastatic tissues (Fig. 3E, Fig. S3A and S3B). Notably, genes such as TMEM201 and RABGAP1L, which are related with cell motility and structural organization exhibited significant upregulation at the isoform level in primary ovarian cancer. In contrast, isoforms of genes such as DUSP22 which is a dual-specificity phosphatase implicated in MAPK signaling modulation, GRPEL2 which is a mitochondrial chaperone showed significantly increased expression levels in metastatic tumors. To validate the results, we experimentally confirmed the isoform switch in TMEM201 in five patient samples using RT-PCR. We selected TMEM201 for experimental validation because it represents a typical example of isoform-level regulation without a corresponding change in total gene expression (Fig. 3F). TMEM201 (also known as NET5) encodes a nuclear envelope transmembrane protein that participates in nuclear positioning and cell migration. It produces two major transcript variants—a short isoform (TMEM201-S) and a long isoform (TMEM201-L). Consistent with the LR-seq results, TMEM201-S showed decreased relative usage in cancer samples, accompanied by a shift toward TMEM201-L (Fig. 3G). This isoform switch is likely to modify the protein’s domain composition, particularly within the DUF2448 and intrinsically disordered regions, thereby influencing nuclear envelope organization and cell motility during tumorigenesis.

### Discovery of alternative splicing events with clinical implications

We analyzed alternative splicing (AS) events from LR-seq data to systematically assess transcript diversity at the splicing level. A total of 22,297 AS events detected across seven major AS types, including 7,219 exons skipping (SE), 558 mutually exclusive exons (MX), 3,370 alternative 5’ splice sites (A5), 3,255 alternative 3’ splice sites (A3), 1,946 retained introns (RI), and 4,936 alternative first (AF) and 1,013 last exons (AL) (Fig. 4A and 4B). Among them, 3,018 AS events (13.54%) were identified in our LR-seq data but absent from GENCODE v45 annotation, revealing the substantial contribution of previously unannotated isoforms in HGSOC-associated splicing. The distribution of AS events did not show significant differences across samples, with SE and AF events being the most common, and a slight increase in RI events observed in metastatic samples (Fig. 4C). We next conducted differential analysis of splicing events between cancer and normal tissues (C vs N) and between metastatic and primary tumors (M vs P). Heatmaps of top 30 representative events showed clear sample-specific splicing patterns, particularly for events such as AF in GNAS or SE in SERPINB6 in cancer samples, and RI or AF alterations in TOP2B, KRAS, and DCAFB in metastatic tumors (Fig. 4D). Functional enrichment analysis of genes exhibiting differential AS events between cancer and normal tissues revealed that distinct AS types are associated with specific biological pathways and processes (Fig. 4E). RI events produced the strongest enrichment signals across multiple categories, including translation, mRNA transport, and RNA localization, suggesting the broad impact of intron retention on translational control. A5 events also contributed to translation-related terms, whereas AF and AL events were associated with transcriptional and genomic-regulatory functions. MX and SE events corresponded to immune signaling and protein-turnover processes. In metastatic versus primary tumors, RI events again exhibited the strongest enrichment, mainly involving translation, RNA splicing, and RNA localization pathways. A5 events were linked to oxidative phosphorylation and oncogenic signaling, whereas AL events were associated with immune diversification and viral-response processes. Overall, intron-retention and splice-site variations dominate translational and RNA-processing regulation, while other splicing modes contribute to cell-cycle, immune pathways during ovarian-cancer progression.

**Fig. 4.**
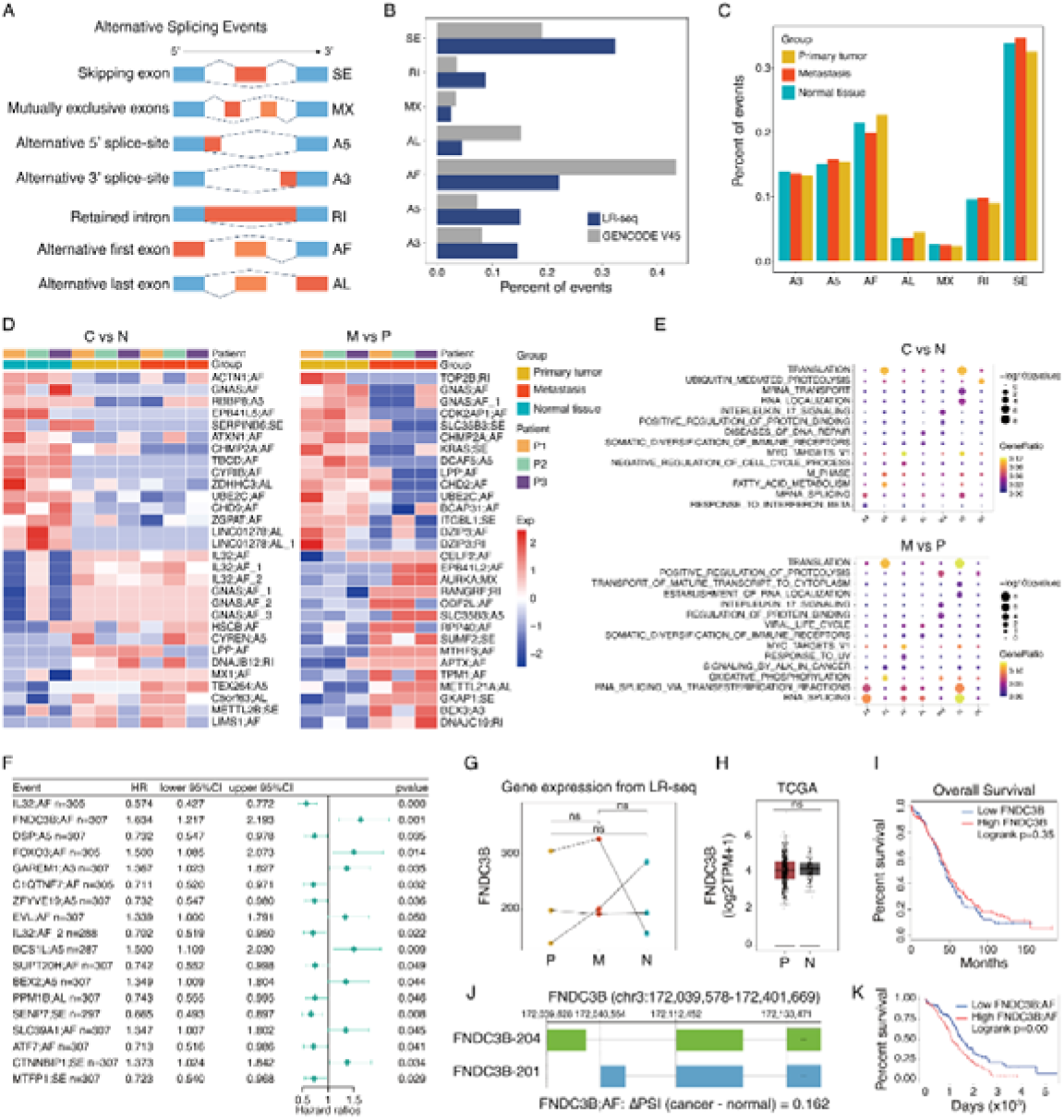
Discovery of alternative splicing events related to HGSOC. (A) Illustration of seven major types of alternative splicing (AS) events, including skipping exon (SE), mutually exclusive exon (MX), alternative 5′ splice-site (A5), alternative 3′ splice-site (A3), retained intron (RI), alternative first exon (AF), and alternative last exon (AL). (B) Comparison of AS event frequencies between LR-seq data and GENCODE v45 annotation. (C) Distribution of AS event types across normal, primary, and metastatic tissues. (D) Top 30 AS events in cancer versus normal (C vs N) and metastatic versus primary (M vs P) comparisons. (E) Enrichment analysis of genes containing differential AS events. (F) Cox proportional hazards regression analysis of AS events. (G–H) Gene-level expression of FNDC3B from LR-seq and TCGA database. (I) Overall survival of high and low FNDC3B gene expression group in TCGA ovarian cancer cohort. (J) Schematic of alternative first-exon (AF) usage in FNDC3B. (K) Overall survival of high and low FNDC3B:AF group.

To evaluate the clinical relevance of these splicing alterations, annotated alternative splicing events were analyzed using the TCGA cohort, and univariate Cox proportional hazards regression analyses were conducted for events that exhibited differential patterns in the long-read sequencing data. Several AS events were significantly associated with overall survival (Fig. 4F). For instance, AF usage in IL32 was associated with better prognosis (HR=0.574, p<0.001), while AF in FNDC3B and SE in CTNNB1 were linked to poor prognosis. FNDC3B was selected for further analysis because its AF event showed the highest hazard ratio among candidates. In both our long-read dataset and the TCGA cohort, total FNDC3B expression levels have no difference between cancer and normal tissues, and the expression was not significantly associated with patient survival (Fig. 4G–H). However, splicing analysis identified AF event of FNDC3B showed changes between cancer and normal samples (Fig. 4J). Patients exhibiting higher inclusion of this cancer-associated isoform displayed significantly poorer overall survival (Fig. 4I–K). These findings demonstrate that gene-level measurements alone can obscure clinically meaningful transcriptomic changes, underscoring the importance of isoform-level analyses for uncovering prognostic alterations that are invisible in conventional expression profiling.

### Discovery of alternative polyadenylation events

To further investigate post-transcriptional regulation beyond splicing, we analyzed alternative polyadenylation (APA) dynamics from our LR-seq data. A total of 30,104 transcription start sites (TSSs) and 27,246 transcription end sites (TESs) were identified from the annotated isoforms. Approximately 70% of the genes contained one or two TSS or TES sites, about 20% had three or four, and the remaining genes possessed more than five TSSs and TESs (Fig. 5A). The majority of TESs and TSSs were located within annotated 3’ and 5’ untranslated regions (UTRs), respectively, followed by exonic regions, and their positional distributions were largely consistent across normal, primary, and metastatic tissues (Fig. 5B–C). Since more than half of the genes used multiple PASs, we next examined whether ovarian cancers exhibit a preference for specific PAS usage. We quantified distal PAS usage (DPU) for each gene, defined as the proportion of transcript counts using distal PAS relative to total counts (Fig. 5D). The DPU value ranges from 0 to 1, where 1 represents exclusive distal PAS usage and 0 indicates exclusive proximal PAS usage. Although the overall DPU distributions were comparable across tissue types, a slight increase of distal PAS usage was observed in metastatic tumors. Differential APA analysis identified multiple genes exhibiting significant 3’-end remodeling between cancer and normal tissues as well as between metastatic and primary tumors (Fig. 5E). Genes such as USP11, CRISPLD2, and BAZ2A showed a preference for distal PAS usage in cancer, whereas PARD3 and RBPJ tended to favor proximal sites. In metastatic versus primary tumors, several genes, including SCAMP1, BIRC6, and DCP2, exhibited a distal shift. Because the alternative choice of TESs is determined by isoform-specific differences and may fail to accurately capture UTR usage, UTR regions were extracted by integrating the existing reference annotation with the previously predicted protein-coding sequences for downstream analyses. We further evaluated 3′UTR length variation by computing the UTR usage ratio (UTRU), defined as the proportion of the longest UTR isoforms among all transcripts (Fig. 5F, Fig. S4A). Similar patterns were observed for 3’UTR length distribution, where metastatic tumors tent to produce longer UTRs. Moreover, numerous genes exhibited significant differential UTR usage (dUTRU) between cancer and normal tissues, and to a lesser extent between metastatic and primary tumors (Fig. 5G). Notably, transcripts such as AKAP12, PRPF4B, and SNX6 had elongated 3′UTRs in cancer, potentially influencing mRNA stability or microRNA binding. These analyses reveal widespread remodeling of 3 ′ -end formation and UTR length in ovarian cancer, suggesting that APA contributes an additional layer of post-transcriptional regulation that distinguishes tumor and metastatic transcriptomes from normal ovarian tissue. Finally, we evaluated the overall changes of post-transcriptional regulation revealed by LR-seq. We calculated the TSS, TES, and splicing ratio scores for each gene in all groups, and visualized the overall splicing patterns using ternary plots. Compared with the GENCODE annotation, the LR-seq data exhibited a stronger bias toward splicing events (Fig. S4B and S4C). However, no significant differences were observed either in the overall splicing patterns or among the three groups when analyzed separately. Subsequently, the ternary plot was divided into four equal parts by the median lines, representing three regions dominated by high TSS, TES, or splicing ratio scores, as well as a mixed region. In addition, the central point of the triangle was defined as representing genes with simple splicing patterns. We counted the number of genes distributed in each region and at the central point, and found that the variation in the splicing-dominated region was the most pronounced across groups (Fig. S4D). Notably, the primary tumor group exhibited greater splicing diversity than the other groups.

**Fig. 5.**
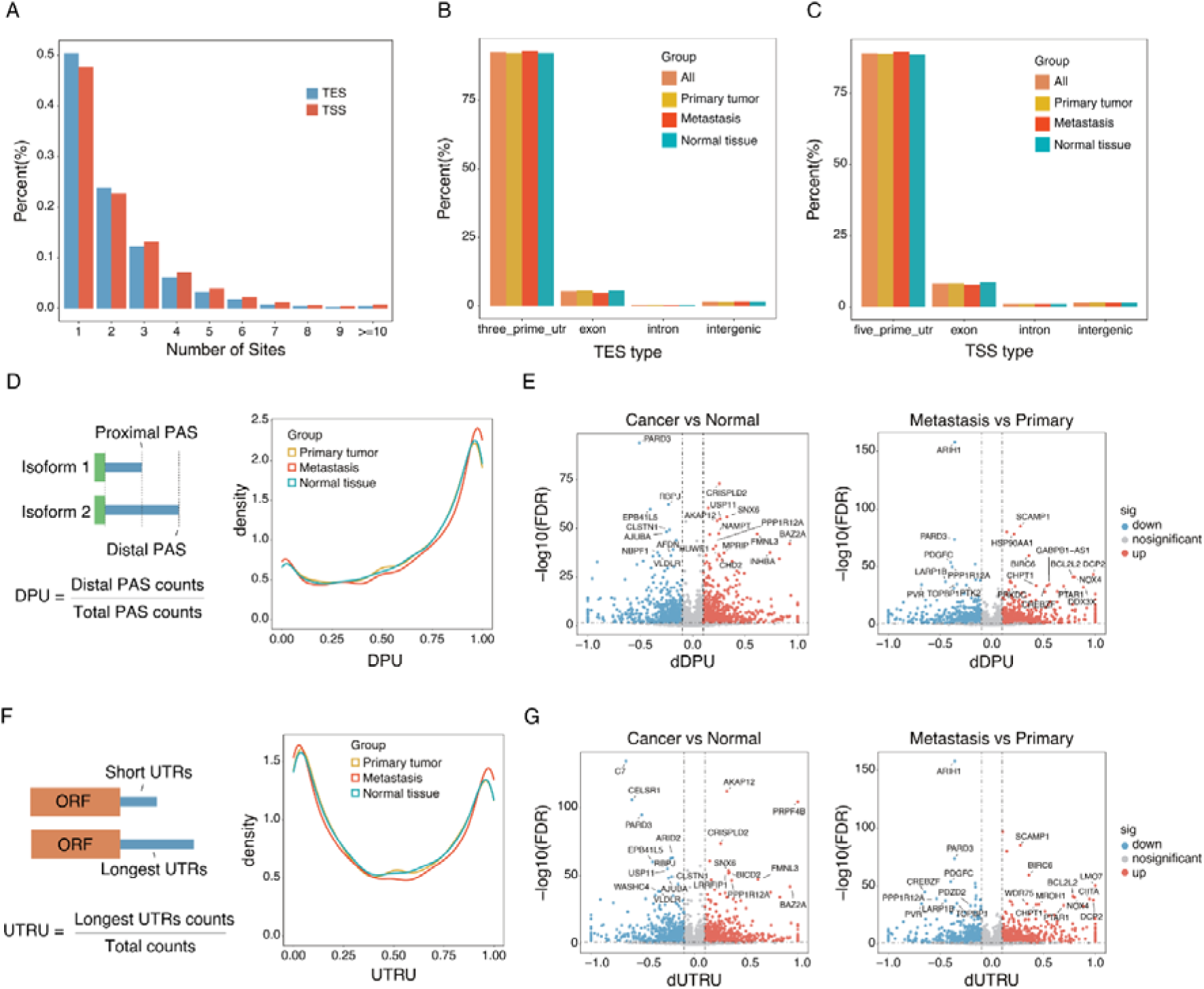
Exploration of alternative polyadenylation in HGSOC. (A) The number of transcription start sites (TSSs) and transcription end sites (TESs) per gene. (B–C) Genomic distribution of TESs and TSSs across tissue groups. (D) Quantification of distal polyadenylation site usage (DPU). The schematic (left) illustrates the definition of DPU as the fraction of distal PAS usage over total PAS usage. Density plots (right) show comparable DPU distributions among groups. (E) Volcano plots of differential APA analysis. (F) Quantification of 3′UTR usage ratio (UTRU). (G) Differential UTR usage (dUTRU) analysis results.

## Discussion

In this study, we performed LR-seq for matched normal, primary, and metastatic tissues from patients with high-grade serous ovarian cancer (HGSOC) to examine complex transcriptome dynamics during cancer progression. We generated a full-length transcriptome map by integrating long- and short-read data, uncovered various previously unannotated isoforms with coding potential, and showed that post-transcriptional remodeling operates at both alternative splicing and polyadenylation level. Recent advances in LR-seq technologies have enabled isoform-resolved analyses across diverse cancer types, such as colorectal[11], gastric[14], cervical[25] and breast cancers[26]. Although a recent single-cell targeted sequencing study has begun to explore isoform diversity in ovarian cancer[27], our work provides a unique perspective by leveraging matched normal, primary, and metastatic samples from the same individuals and by presenting, to the best of our knowledge, the first comprehensive long-read transcriptomic analysis of metastatic ovarian tumors.

During analyses, we observed that gene-level changes capture only part of the biology, whereas isoform-level alterations reveal additional mechanisms relevant to tumor initiation and progression. Although many genes were differentially expressed between conditions, only a small fraction showed concurrent DTU, indicating that qualitative remodeling of transcript structures can occur independently of total abundance. This decoupling demonstrates why isoform-resolved analyses are necessary to interpret tumor biology beyond bulk expression. The isoform switch analyses provide typical examples of such qualitative control. We detected several switches that alter open reading frames, intrinsically disordered regions, protein domains, susceptibility to nonsense-mediated decay, and predicted subcellular localization. Notably, TMEM201 illustrated an isoform switch without a corresponding change in total gene expression; validation across independent samples confirmed reduced usage of the short isoform and increased expression of the long isoform in tumors. These findings suggest that HGSOC may alter protein features and cellular behavior through changes in isoform composition rather than simply up- or down-regulating genes.

We explored the prognostic relevance of alternative splicing in ovarian cancer using the TCGA cohort. Among the AS events identified from LR-seq data, 18 of these events showed significant associations with overall survival. These results indicate that although splicing alterations are widespread in cancer transcriptomes, only a limited fraction appear to influence patient outcomes, highlighting their potential as selective prognostic biomarkers. Of the genes associated with prognostic splicing events, several genes have previously been implicated in cancer progression. For example, IL32, a secreted protein containing an RDG domain, is highly expressed in cancer-associated fibroblasts and promotes breast cancer cell invasiveness through direct binding[28]. FOXO3, a member of the forkhead box (FOX) transcription factor family, binds to the SIRT6 promoter region upon dephosphorylation, triggering *SIRT6* transcription and inducing mitochondrial apoptosis[29]. PPM1B functions as a negative regulator of the NF-κB signaling pathway and suppresses the migration and invasion of colorectal cancer cells[30]. Furthermore, the analysis also uncovered an alternative first-exon events of FNDC3B were associated with patients’ survival. In parallel, analysis of 3′-end processing identified extensive APA activity across ovarian cancer samples. Earlier studies have reported that malignant cells exhibit global 3′UTRs shortening, thereby escaping microRNA- or RNA-binding–protein– mediated repression[31, 32]. However, with the advent of high-throughput technologies, emerging evidence suggests that 3′UTR lengthening can also occur in specific cancers and is often associated with poor clinical outcomes[33, 34]. Our data also showed no widespread shortening in cancer compared with normal tissues; instead, metastatic tumors exhibited a tendency toward longer 3′UTRs. Lengthened 3′UTRs can expand regulatory potential by introducing additional microRNA binding sites and modulating mRNA localization, suggesting that metastatic cells may exploit UTR extension to fine-tune gene expression during dissemination. Further validation in larger patient cohorts will be required to confirm whether UTR lengthening consistently correlates with metastatic potential and clinical outcomes.

This work has some limitations. The cohort size is small (three patients with matched tissues), which constrains power to detect rare or patient-specific events and may limit generalizability. Moreover, tissue heterogeneity can influence isoform measurements, and although long-read sequencing reduces assembly ambiguities, platform-specific biases and depth differences relative to short-read data remain. Future studies should expand to larger, clinically annotated cohorts with orthogonal biological validation and incorporate single-cell or spatial transcriptomics to capture cell-type–specific isoform programs.

In summary, our long-read transcriptome analysis highlights multilayered post-transcriptional reprogramming in HGSOC and shows that isoform-level information yields biological and clinical insights not accessible from gene-level profiling alone. These results motivate an isoform-aware view of ovarian cancer biology and open avenues for biomarker development and targeted intervention at the level of RNA processing.

## Methods

### Paired normal tissue and ovarian cancer specimens

Paired normal ovarian tissues and ovarian carcinoma samples were obtained. The study was approved by the Ethics Committee of xxx University. All participants provided written informed consent, and the research followed the principles of the Declaration of Helsinki.

### Transcriptome sequencing

Fresh ovarian cancer tissue samples were snap-frozen in liquid nitrogen and ground. Total RNA was extracted from normal ovarian tissue, primary tumor tissue, and metastatic lesions of ovarian cancer patients using TRIeasyTM Total RNA Extraction Reagent according to the manufacturer’s instruction.

For long-read RNA sequencing, polyadenylated RNA molecules were enriched with Oligo(dT) primers and reverse-transcribed to synthesize cDNA Using the Iso-Seq Express 2.0 kit, followed by PCR amplification to generate barcoded cDNA. The Kinnex Full-Length RNA kit was then used to amplify the cDNA and introduce sticky ends, allowing concatenation of eight cDNA fragments in series while ligating barcoded adapters at both ends. Damage repair was performed, and exonuclease treatment was applied to remove incompletely ligated fragments, yielding the final sequencing library. After quantification, the library templates of a certain concentration and volume and enzyme complexes were loaded into the nanopores of the PacBio Revio sequencing system for single-molecule real-time (SMRT) sequencing.

For short-read RNA sequencing, eukaryotic mRNA was enriched using Oligo(dT) magnetic beads. The enriched mRNA was then fragmented using fragmentation buffer. Using the fragmented mRNA as a template, first- and second-strand cDNA were synthesized and subsequently purified. The purified double-stranded cDNA underwent end repair, A-tailing, adapter ligation, and fragment size selection to obtain cDNA fragments of approximately 350 bp. Finally, PCR amplification was performed to generate the cDNA library. The double-stranded cDNA was then denatured at high temperature to produce single-stranded DNA, followed by the addition of a circularization primer to form a single-stranded circular library. The constructed libraries were quality-checked using Qubit 3.0 and Agilent 2100 systems. Libraries that passed quality assessment were sequenced on a high-throughput sequencing platform with a PE150 strategy.

### Iso-seq long-read processing

Raw PacBio HiFi reads were produced on the Revio system and already removed primers. The pipeline of data processing was obtained from https://isoseq.how/. Two major steps were performed to obtain full-length non-chimeric (FLNC) reads: (i) Poly(A) tails and concatemers were identified and removed to refine high-quality full-length (FL) reads; (ii) The resulting FL reads from multiple SMRT cells of the same sample were merged and clustered to generate high-confidence transcript isoforms. The processed FLNC reads were then aligned to the human reference genome (hg38) using pbmm2 with the ISOSEQ preset. Aligned BAM files were subsequently sorted and indexed with Samtools.

### RNA-seq short-read processing

Raw sequencing reads were mapped to human genomes (hg38) using STAR[35]. The resulting alignments were subsequently sorted and indexed with Samtools. Gene-level counts were then obtained using featureCounts.

### Isoform quantification and annotation

Sorted bam files from long read were quantified using isoquant based on hg38 genomes and GENCODE comprehensive v45 annotations. Sorted bam files from short read were quantified using StringTie[36]. Genes and transcripts detected annotations and quantification counts were generated. Detected unique isoforms were annotated using SQANTI3[37] with GENCODE comprehensive v45. Isoforms were characterized and categorized into structural categories.

### Protein-level analysis

Coding regions of LR-seq–derived isoforms were evaluated for homology and domain conservation against the human proteome reference in UniProt. UniProt release 2024-03 (Swiss-Prot) and VarSplice was used as reference, which contains 571,609 protein sequences. TransDecoder (version 5.7.1) was adopted to predict coding regions from transcript annotation files as described below. ORFs with a minimum length of 100 amino acids were retained for downstream analysis. Diamond blastp (version 0.8.36) was used to align ORFs to the reference proteome with parameters max-target-seqs = 1 and evalue = 1e-5 to identify homologs to UniProt. We also search the extracted ORFs based on Pfam database for protein domains using hmmsearch from hmmer (version 3.1b2) with default parameters. Coding regions prediction was performed then integrating the UniProt homology and domain conservation to human proteins to determine the confident protein-coding regions. Global alignment was performed to calculate the identity between transcript coding regions and homologs in Uniprot using Biostrings (version 2.66.0) in R. TMHMM (version 2.0) was used to predict transmembrane helices[38]. CPC2[39] was applied to predict the coding potential of isoforms, IUPred2A[40] was employed to predict intrinsically disordered regions and disordered binding regions within protein sequences, DeepLoc2 was used to predict subcellular localization, and SignalP 6.0 was applied to identify signal peptide regions. The code in this section referenced previous paper[26].

### DGE, DTE, and DTU

EdgeR (version 3.40.2) was employed to perform differential analysis. To detect changes in relative transcript abundance, DRIMseq (version 1.30.0)[41] was used for differential transcript usage analysis. All packages above were created in R script (version 4.3.2). The overall analysis of alternative isoform usage and the identification of isoform switching events were performed using IsoformSwitchAnalyzeR[42].

### Splicing analysis

AS events were determined and quantified using SUPPA2 (version 2.4), based on GTF files generated by SQANTI3 while isoform annotation. SUPPA2[43] classifies AS events into alternative 3′ splicing (A3), 5′ splicing (A5), first exon (AF), last exon (AL), intron retention (RI), exon skipping (SE) and mutually exclusive exons (MX). The transcript expression files were produced by isoform quantification and normalized to TPM (transcripts per million) values. AS inclusion levels (PSI) were estimated from transcript quantifications using SUPPA2, and statistical significance between conditions was assessed with the Kolmogorov–Smirnov test to identify high-confidence AS events.

Transcript structural features were analyzed using Cerberus, which summarized transcriptional diversity by quantifying the usage of TSSs, exon junction chains (ECs), and TESs for each. Splicing ratio was calculated as 2 × (number of ECs) / (number of TSSs + number of TESs) to properly represent the contribution of ECs to transcript diversity. Using TSS ratio, splicing ratio and TES ratio, Cerberus categorized genes into TSS-high, splicing-high, TES-high, mixed for genes with multiple transcripts that do not prefer each mode and simple for genes with just one transcript. A simplex plot on three variables was used to visualize the preference of each group.

### TCGA analysis

TCGA isoform-level expression matrix, clinical information, and survival data were downloaded from the Firehose repository. The isoform identifiers in the expression matrix were first converted from kg5 to kg9 IDs, and finally to Ensembl IDs to ensure consistency with downstream analyses. AS events were determined and quantified using SUPPA2, based on GTF files from GENCODE v45. Survival analyses for TCGA samples were conducted using the survminer and survival packages in R.

### TSS and TES analyses

The transcription start sites (TSS) and transcription termination sites (TTS) were derived from the annotated isoforms. To identify alternative polyadenylation events, the most distal 3’ end point among all annotated transcripts for each gene was defined as distal poly(A) site, all the other poly(A) sites were distinguished as proximal poly(A) site. Distal poly(A) site usage (DPU) was used to measure poly(A) site usage of each gene among all groups. We calculated absolute difference of DPU (dDPU) between conditions. The annotation of untranslated regions (UTRs) was derived by integrating GENCODE v45 annotation with the previously predicted protein-coding regions. For each gene, the isoform harboring the longest UTR was identified as the long-UTR isoform, whereas all others were classified as short-UTR isoforms. 3 ′ UTR usage was estimated by calculating the fraction of transcripts carrying the long-UTR isoform relative to the total isoform expression of each gene. Group-level differences in UTR usage were evaluated. All statistical significance in this part was determined using Fisher’s exact test, with P-values adjusted for multiple testing by the Benjamini–Hochberg procedure. All the above analyses were conducted allowing a 10 bp tolerance.

### Enrichment analysis

Gene sets used in the article are all from human MSigDB collections. Over-representation analysis was performed by enricher function from clusterProfiler (version 4.10.0) in R.

### PCR/RT-qPCR

cDNA was synthesized from 1 μg of RNA using using M-MLV reverse transcriptase (Promega, Madison, WI). Quantitative real-time PCR was performed using SYBR qPCR Master Mix (TransGen Biotech, Beijin, China). The PCR was performed using KOD polymerase (TOYOBO). The sequences of the primers used are as follows:

TMEM201-All isoforms-Forward Primer: 5′-CAGATGTACAAGCTGTGCCG-3′;

TMEM201-All isoforms-Reverse Primer: 5′-CGCGGTGGTCAGTAGGAAG-3′;

TMEM201-S -Forward Primer: 5′- GCTCAGCCACACCTGACAATG-3′;

TMEM201-S-Reverse Primer: 5′- GGTGAAAAGAGGCAGGACAGAG-3′;

KRAS Forward: 5′-ACAGGCTCAGGACTTAGCAA-3′;

KRAS Reverse: 5′-TAACAGTCTGCATGGAGCAGG-3′.

## Author contributions

Yongjun Piao and Yi Shi designed the research. Wei Ding provided patient tissues. Tianxiang Liu analyzed data. Jianying Lv and Shuo Wang performed the experiments. Yongjun Piao, Tianxiang Liu, Jianying Lv and Shuo Wang wrote the manuscript. Yanhua Liu, Yanan Chen, Jia Li, Longlong Wang, Wei Ding and Yi Shi supported the design of experimental procedures and were responsible for manuscript review and editing.

## Data availability

All sequencing data can be obtained at Genome Sequence Archive (GSA) under accession number HRA014484.

---

**Fig. S1.**
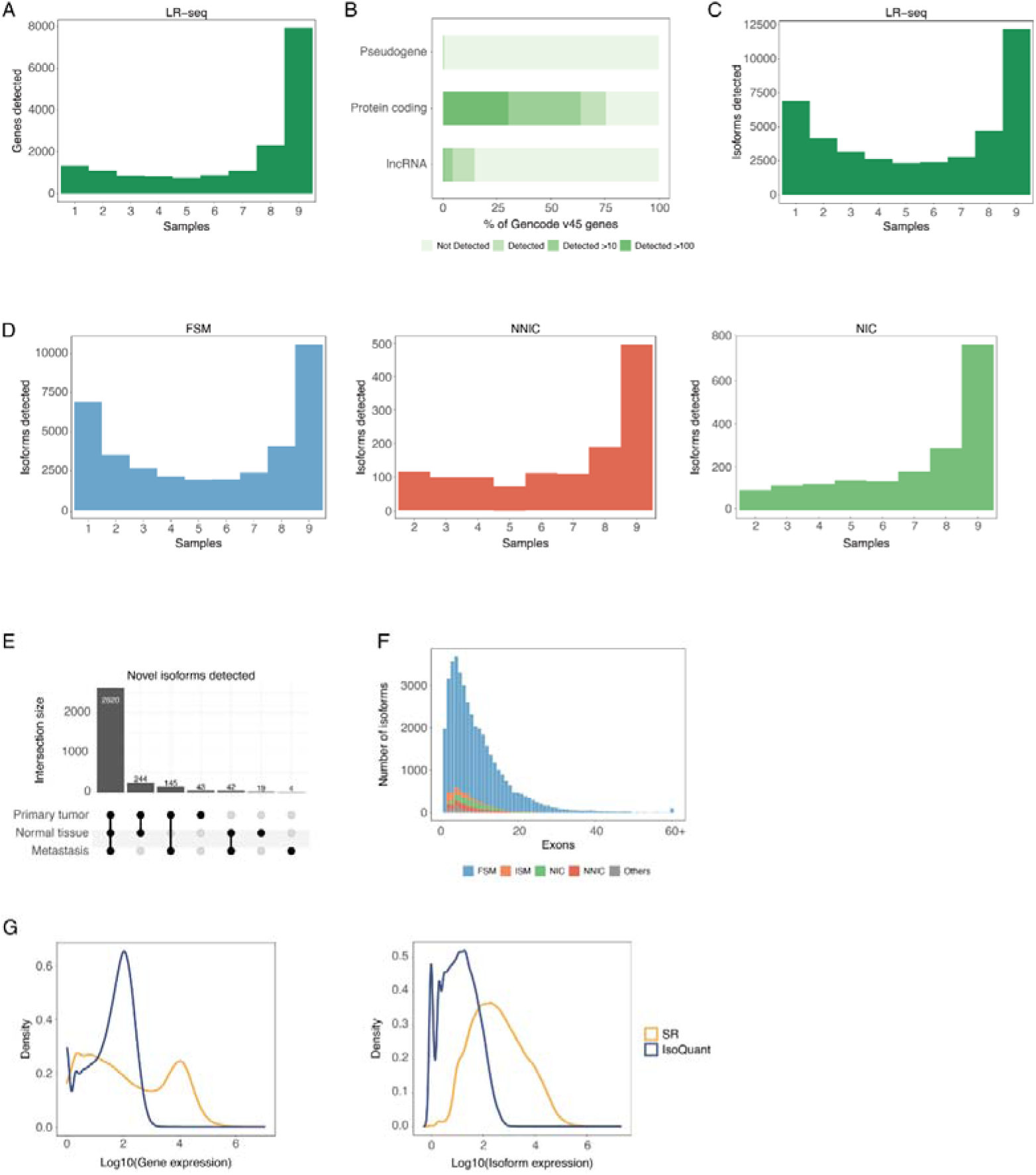
Distribution of detected genes and isoforms among samples. (A) The number of samples in which genes were detected. (B) The proportion of Gencode v45 genes of each biotype detected in samples with TPM > 0, TPM > 10, and TPM > 100. (C) The number of samples in which isoforms were detected. (D) separated by isoform categories. (E) The numbers of novel isoforms (NIC and NNIC) detected in each group. (F) Number of exons distribution for each transcript category. (G) Overall expression distributions between LR-seq and SR-seq at the gene and transcript levels.

**Fig. S2.**
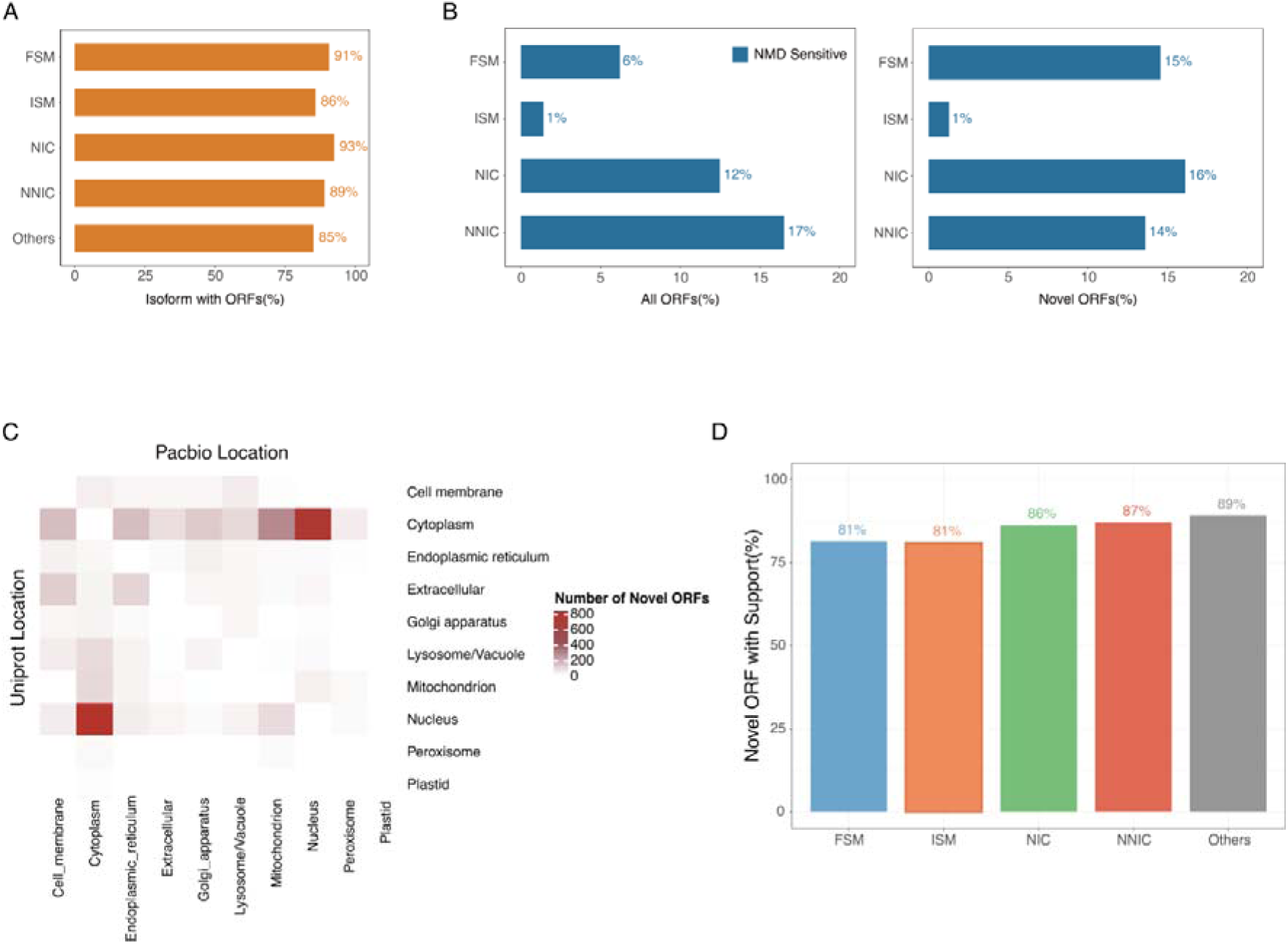
Coding potential and NMD prediction of ORFs from LR-seq isoforms. (A) The proportion of isoforms predicted to contain open reading frames in each category. (B) The proportion of isoforms identified as NMD-sensitive per category for all ORFs or for novel ORFs. (C) Predicted subcellular localization changes of ORFs from LR-seq isoforms compared to the most similar protein in UniProt. (D) Proportion of novel ORFs verified by MS/MS proteomics per category.

**Fig. S3.**
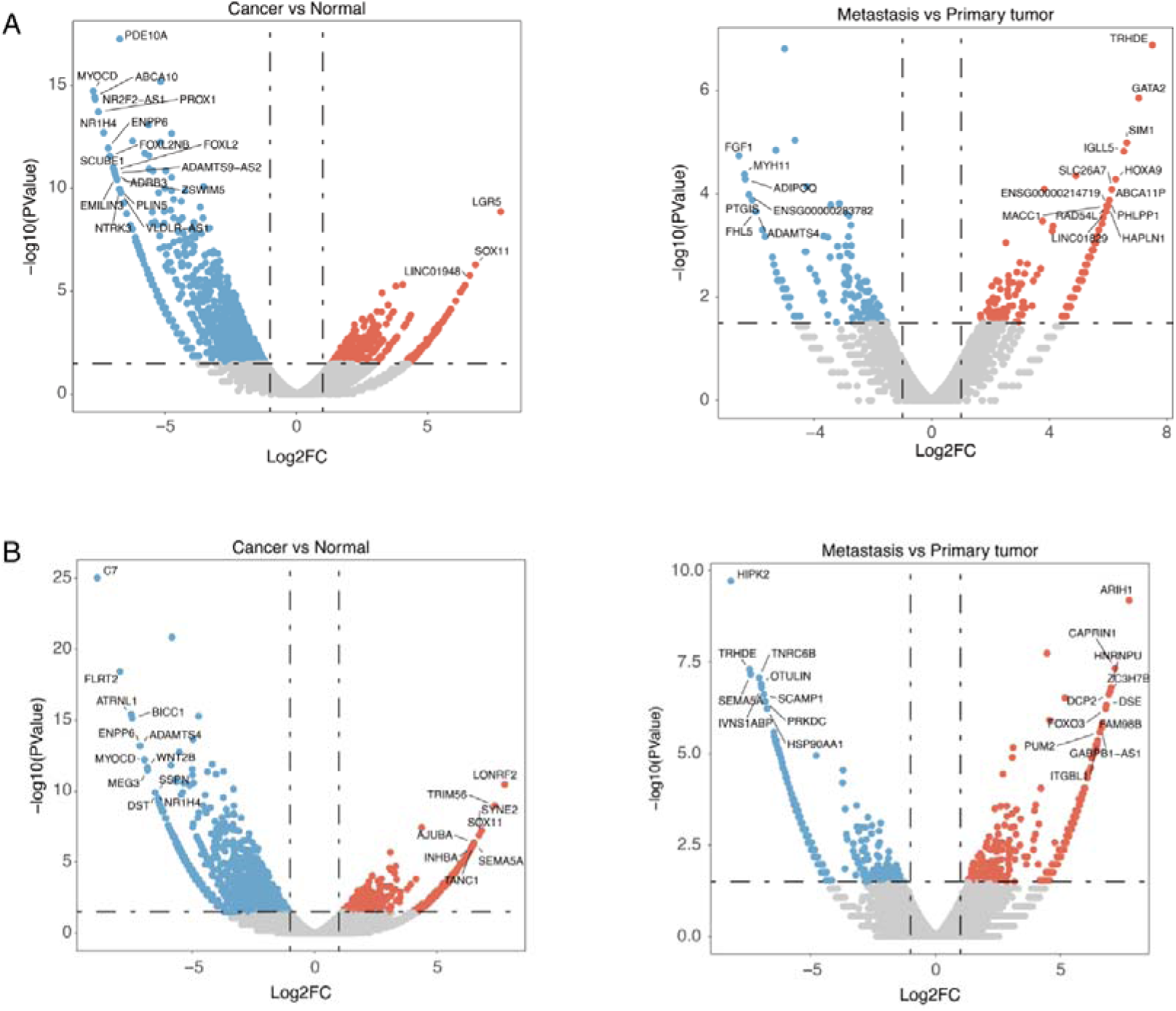
Volcano plots for DGE and DTE. (A) Volcano plots of genes with significant DGE between cancer and normal samples (left) as well as between metastatic and primary tumors (right). (B) Volcano plots of genes with significant DTE between cancer and normal samples (left) as well as between metastatic and primary tumors (right).

**Fig. S4.**
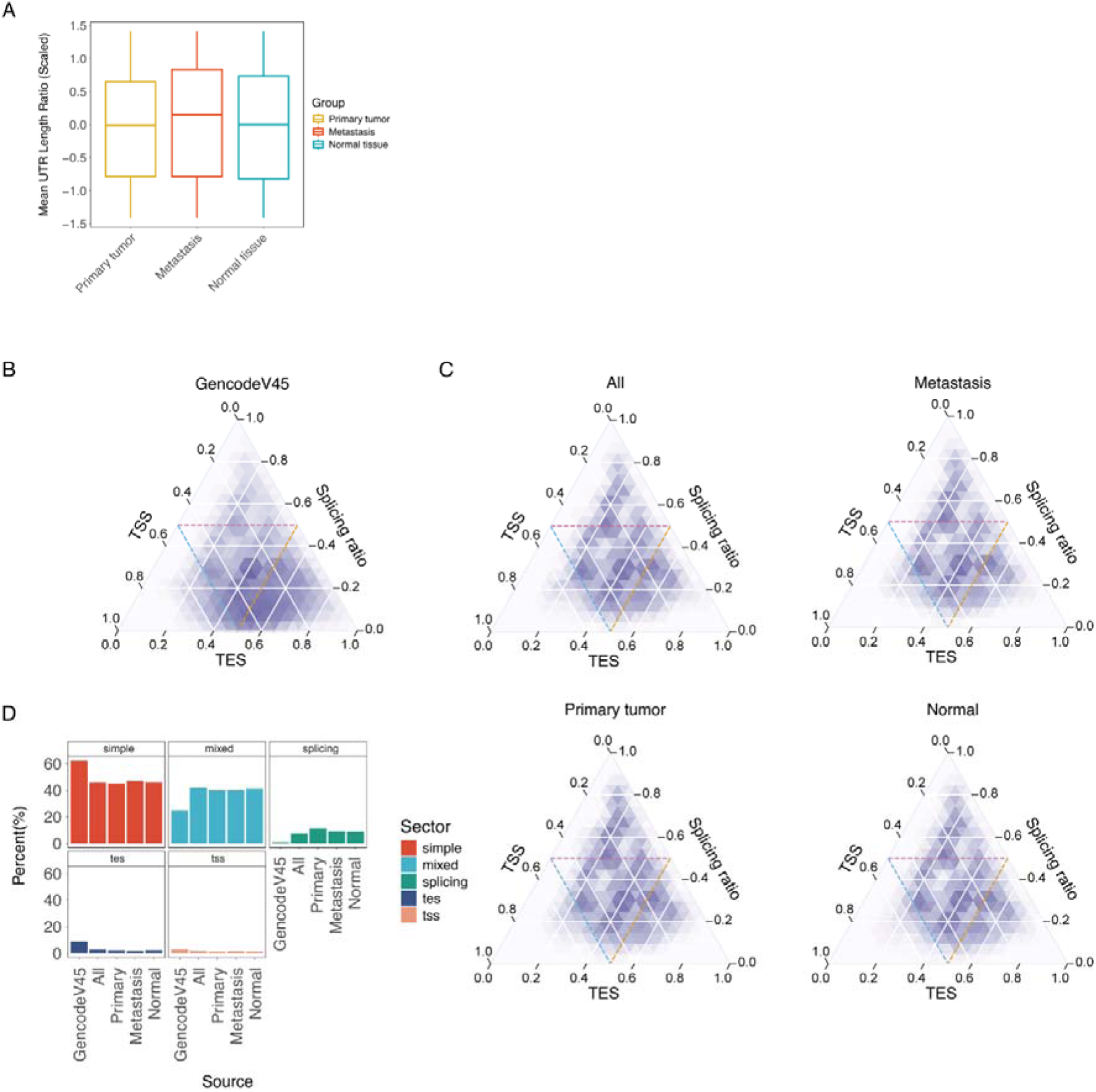
The overall changes of post-transcriptional regulation revealed by LR-seq. (A) Mean UTR length ratio of genes in each groups. (B) Overall splice pattern distribution in Gencode45. (C) Overall splice pattern distribution in all samples and each group. (D) Proportion of genes located in each region in (B) and (C).

**Table S1. Basic information of sequencing files in the raw and processing workflow.**

**Table S2. The results of the difference analysis, including DGE, DTE and DTU.**

